# *Drosophila* insulin-like peptide 8 (DILP8) in ovarian follicle cells regulates ovulation and metabolism

**DOI:** 10.1101/2020.05.02.073585

**Authors:** Sifang Liao, Dick R. Nässel

## Abstract

In *Drosophila* eight insulin-like peptides (DILP1-8) are encoded on separate genes. These DILPs are characterized by unique spatial and temporal expression patterns during the lifecycle. Whereas functions of several of the DILPs have been extensively investigated at different developmental stages, the role of DILP8 signaling is primarily known from larvae and pupae where it couples organ growth and developmental transitions. In adult female flies, a study showed that a specific set of neurons that express the DILP8 receptor, Lgr3, is involved in regulation of reproductive behavior. Here, we further investigated the expression of *dilp8*/DILP8 and *Lgr3* in adult female flies and the functional role of DILP8 signaling. The only site where we found both *dilp8* expression and DILP8 immunolabeling was in follicle cells of mature ovaries. *Lgr3* expression was detected in numerous neurons in the brain and ventral nerve cord, a small set of peripheral neurons innervating the abdominal heart, as well as in a set of follicle cells close to the oviduct. Ovulation was affected in *dilp8* mutants as well as after *dilp8*-RNAi using *dilp8* and follicle cell Gal4 drivers. More eggs were retained in the ovaries and fewer were laid, indicating that DILP8 is important for ovulation. Our data suggest that DILP8 signals locally to *Lgr3* expressing follicle cells as well as systemically to *Lgr3* expressing efferent neurons in abdominal ganglia that innervate oviduct muscle. Thus, DILP8 may act at two targets to regulate ovulation: follicle cell rupture and oviduct contractions. Furthermore, we could show that manipulations of *dilp8* expression affect food intake and starvation resistance. Possibly this reflects a feedback signaling between ovaries and the CNS that ensures nutrients for ovary development. In summary, it seems that DILP8 signaling in regulation of reproduction is an ancient function, conserved in relaxin signaling in mammals.

## Introduction

Eight insulin-like peptides (DILP1-8), encoded on separate genes, have been identified in *Drosophila* (Brogiolo et al., 2001;Grönke et al., 2010;Colombani et al., 2012;Garelli et al., 2012;Nässel and Vanden Broeck, 2016;Ahmad et al., 2019). These are characterized by unique spatial and temporal expression patterns during the lifecycle [reviewed in (Brogiolo et al., 2001;Owusu-Ansah and Perrimon, 2014;Nässel and Vanden Broeck, 2016;Ahmad et al., 2019)]. Several of the DILPs, notably DILP2, 3, 5 and 6, have been extensively investigated in their roles in development, reproduction, metabolism and effects on lifespan [see (Rulifson et al., 2002;Broughton et al., 2005;Okamoto et al., 2009;Slaidina et al., 2009;Owusu-Ansah and Perrimon, 2014;Nässel and Vanden Broeck, 2016)]. The most recently discovered one, DILP8, acts on a relaxin receptor homolog Lgr3, which is a leucine-rich repeat containing G-protein coupled receptor (GPCR) (Colombani et al., 2012;Garelli et al., 2012;Colombani et al., 2015;Garelli et al., 2015;Jaszczak et al., 2016;Gontijo and Garelli, 2018). DILP8/Lgr3 have been studied in the larva where they are part of a circuit that regulates timing of development of adult progenitor cells in imaginal discs by controlling production of the steroid hormone ecdysone (Ecd) (Colombani et al., 2012;Garelli et al., 2012;Colombani et al., 2015;Garelli et al., 2015;Vallejo et al., 2015;Jaszczak et al., 2016;Juarez-Carreño et al., 2018). DILP8 is produced in imaginal discs upon damage or tumor development and is released to act on Lgr3 in a small set of brain neurons that in turn inhibit release of prothorcicotropic hormone (PTTH) by acting on four lateral neurosecretory cells that innervate the prothoracic gland. This results in diminished production of Ecd and 20-hydroxy-Ecd (20E) (Colombani et al., 2012;Garelli et al., 2012;Colombani et al., 2015;Garelli et al., 2015;Vallejo et al., 2015;Jaszczak et al., 2016;Juarez-Carreño et al., 2018). Thus, these studies show that when discs are damaged DILP8 signaling ensures that metamorphosis is delayed and growth of imaginal discs is slowed down, allowing regeneration of imaginal discs and symmetric growth of the adult organism. A role of *dilp8* in interorgan signaling has also been shown in early pupae upon increased Ras/JNK signaling (Ray and Lakhotia, 2019). However, adult roles of DILP8 signaling are poorly known.

In adult female flies *dilp8* transcript seems to be primarily expressed in the ovaries and very low levels were detected in tissues of male flies (Gontijo and Garelli, 2018) [see also FlyAtlas, Flyatlas.org (Chintapalli et al., 2007) and FlyAtlas2, http://flyatlas.gla.ac.uk/FlyAtlas2/index.html, (Leader et al., 2017)]. Also the *Lrg3* expression is sexually dimorphic in the adult CNS (Meissner et al., 2016). To our knowledge, only one study has explored the functional role of the DILP8/Lgr3 signaling system in adult flies. Thermogenetic activation of a subset of *Lgr3* expressing neurons in the abdominal ganglion reduced receptivity in reproductive behavior of females and decreased fecundity (Meissner et al., 2016), suggesting a relaxin-like function of DILP8 in flies.

Here we undertook a study of the role of DILP8/Lgr3 in adult physiology, including fecundity. Utilizing *dilp8* mutant flies as well as the Gal4-UAS system (Brand and Perrimon, 1993) to manipulate levels of *dilp8* in a targeted fashion, we confirm a role in ovary function and ovulation, but also see effects on feeding and starvation resistance. We find that *dilp8*/DILP8 is expressed in follicle cells in egg chambers of ovaries in young flies, whereas *Lgr3* expression is primarily in neurons of the central and peripheral nervous systems, some of which are efferents that innervate oviduct muscle. Furthermore, a small number of follicle cells in the basal region of each egg chamber express the receptor. Thus, DILP8 participates in both interorgan and intraorgan signaling. The peptide seems to signal systemically from the ovaries to efferent neurons that in turn regulate follicle cell function and affect ovulation and fecundity. In addition, DILP8 acts more locally, in a paracrine fashion, on small sets of basal follicle cells. Taken together, our data support an evolutionarily conserved function of relaxin-like peptides in ovulation and fecundity.

## Materials and Methods

### Fly stocks and maintenance

The following fly strains were used in this study: *w^1118^*, Canton S, UAS-mCD8-GFP, w*, P{w[+mC]=UAS-GFP.S65T}, *Elav* gene-switch-Gal4 (Elav^*GS*^-*Gal4*) and w, lexAop-CD2 RFP; UAS-CD4-spGFP1-10, lexAop-CD4-spGFP11 were obtained from Bloomington *Drosophila* Stock Center (BDSC), Bloomington, IN, USA. Four *Lgr3*-Gal4 lines, *Lgr3*-Gal4∷VP16 (III), *Lgr3*-Gal4∷VP16/cyo, R19B09-Gal4, *Lgr3*-Gal4∷p65/TM3,Sb, and a UAS-line, JFRC81-10XUAS-Syn21-IVS-GFP-p10 (Pfeiffer et al., 2008;Meissner et al., 2016), were obtained from Michael Texada (Janelia Farm, Ashburn, VA, USA). An *engrailed* (*en*)-Gal4 (Tabata et al., 1995) was provided by Dr. Vasilios Tsarohas (Stockholm, Sweden); *dilp8^MI00727^* (*dilp8* mutant with enhanced green fluorescent protein (eGFP) trap in the gene’s first intron [Mi{MIC}CG14059MI00727]), *dilp8*-Gal4 and UAS-*dilp8*∷3xFLAG (Garelli et al., 2012) were from Maria Dominguez (Alicante, Spain); *dilp2*-Gal4 (Rulifson et al., 2002) from E. Rulifson (Stanford, CA); *ppl*-Gal4 (Zinke et al., 1999) from M.J. Pankratz (Bonn, Germany), *c929*-Gal4 (Hewes et al., 2003) from Paul H. Taghert (St Louis, MO); and UAS-*dilp8*/TM3,Sb (Colombani et al., 2012) was provided by P. Leopold (Nice, France) and w^1118^;daughterless-GeneSwitch-Gal4 (Tricoire et al., 2009) was from V. Monnier (Gif-sur-Yvette, France). A *dilp2*-lexA∷VP16 (Li and Gong, 2015) was obtained from Zhefeng Gong (Hangzhou, China), forwarded by Annick Sawala and Alex Gould (London, UK). A double balancer Sco/Cyo,dfd-GFP;Dr/TM6bTb,dfd-GFP was used to combine dilp2-lexA∷VP16 with R19B09-Gal4 for GRASP analysis.

Fly stocks were reared and maintained on standard Bloomington medium (https://bdsc.indiana.edu/information/recipes/bloomfood.html) at 18 °C. The experimental flies were kept at 25 °C on an agar-based diet with 10% sugar and 5% dry yeast. To activate *Elav^GS^*-Gal4 in adult stage, flies were fed RU486 (mifepristone; Sigma, St. Louis, MO, USA) dissolved in ethanol and added to the food at a final concentration of 20 μM. The same amount of ethanol was added to the control food.

### Production of DILP8 antisera

To generate of DILP8 antisera we selected two sequences from the *D. melanogaster* DILP8 protein for custom synthesis:

DmDILP8_42-60_: NH_2_-CEHLFQADEGARRDRRSIE-CONH_2_ (19 aa)

DmDILP8_70-84_: NH_2_-CGSGKTHNKHHYISRS-CONH_2_ (16 aa)

These were coupled to thyroglobulin at the N-terminal cysteine (C), and the two peptide conjugates were mixed and injected as a cocktail into two rabbits for three months of immunization. The production of the antigens and the immunization were performed by Pineda Antibody Service (Berlin, Germany). The antisera were affinity purified before use. Preimmune sera did not immunolabel follicle cells when applied at 1:10,000.

### Immunocytochemistry and imaging

Immunohistochemistry of *Drosophila* larval and adult tissues were performed as in previous study (Liu et al., 2016). Tissues were dissected in phosphate-buffered saline (PBS) and then fixed in 4% ice-cold paraformaldehyde (PFA) (2 hours for larval samples and 4 hours for adults), and subsequently rinsed in PBS for 1 h. Samples were then incubated for 48 hours at 4°C in primary antibodies diluted in PBS with 0.5% Triton X-100 (PBST). After washes in PBST for 1 hour at room temperature, the samples were incubated with secondary antiserum for 48 hours at 4°C. Finally, all samples were washed with PBST and PBS, and then mounted in 80% glycerol with 0.1 M PBS. Images were captured with a Zeiss LSM 780 confocal microscope (Jena, Germany) using 10×, 20×, or 40× oil immersion objectives. Images of the whole fly were captured with a Zeiss Axioplan 2 microscope after freezing the flies at −20°C. Confocal and microscope images were processed Fiji (https://imagej.nih.gov/ij/) and for contrast and intensity in Adobe Photoshop.

The following primary antisera were used: rabbit anti-DILP8 (this study) used at 1:10,000; mouse anti-green fluorescent protein (GFP) at 1:000 (RRID: AB_221568, Invitrogen, Carlsbad, CA). Rabbit anti-leucokinin (Nässel et al., 1992) at 1:2000; rabbit anti-ion transport peptide (Galikova et al., 2018) at 1:1500; rabbit anti-DILP3 (Veenstra et al., 2008) at 1:2000 provided by J. A. Veenstra, Bordeaux, France, rabbit anti-tyrosine decarboxylase-2 (Tdc2; pab0822-P, Covalab, Cambridge, UK; at 1:200 [see (Pauls et al., 2018)] obtained from D. Pauls, Leipzig, Germany; Rhodamine-phalloidin was used at 1:1000 to stain muscle. The following secondary antisera were used: goat anti-rabbit Alexa 546, goat anti-rabbit Alexa 488 and goat anti-mouse Alexa 488 (all from Invitrogen). DAPI (Sigma) at a dilution of 1:2000 was used to staining the nuclei.

### Stress resistance experiments

6–7 days old female flies were used for starvation resistance experiments. The flies were placed in vials containing 0.5% aqueous agarose for starvation in an incubator at 25 °C with 12:12 h Light:Dark (LD) conditions and controlled humidity. Dead flies were counted at least every 12 hours. For desiccation resistance the experiment was the same, except that flies were kept in empty vials, thus the flies obtained no food and no water.

### Determination of eclosion time

To determine the eclosion time, one-week-old adult parental flies were mated in the evening. The next morning these mated flies were transferred to vials with fresh food for egg laying during 4 h. After that the adult flies were removed. Two hours after the initiation of oviposition was considered time “0”, and thereafter, the number of flies eclosed from these vials were monitored at least every 12 hours.

### Capillary feeding assay

To study the food intake of individual flies, capillary feeding (CAFE) assay was used according to (Ja et al., 2007). Each fly was kept in a 1.5 mL Eppendorf tube inserted with a 5 μL capillary tube. The capillary tubes were loaded with food containing 5% sucrose, 2% yeast extract, and 0.1% propionic acid. Food consumption was measured every day and the cumulative food intake was calculated and used for generating the graphs. Three biological replicates with 10 flies in each were used for each genotype.

### Oviposition and egg numbers in ovaries of the flies

For oviposition, 5 days old flies were used. The number of eggs laid by individual pairs of flies was counted every 24 hours for two days, and the total number of eggs laid in 48 hours was calculated. After the egg laying experiments, the same flies were used to monitor the number of eggs that are at stage 10 - 14 [according to (Saunders et al., 1989;Shimada et al., 2011;Kubrak et al., 2014)] inside each female fly.

### Statistical Analysis

Prism version 6.00 (La Jolla, CA, USA) was used for statistics and generating the graphs. The experimental data are presented as means ± SEM. Data were checked firstly with Shapiro-Wilk normality test, then one-way analysis of variance (ANOVA) was used for comparisons among three groups or Student’s t-test was performed followed with Tukey’s multiple comparisons test when comparing two groups. Survival data were compared with survival analysis (Log Rank comparison with Mantel-Cox post test).

## Results

### Distribution of dilp8/DILP8

The distribution of DILP8 producing cells was monitored with the Gal4-UAS system (Brand and Perrimon, 1993) using a *dilp8*-Gal4 line to drive GFP (Fig. 1A). We also employed immunocytochemistry with a novel antiserum to DILP8 (Fig. 1B) and a *dilp8* mutant with enhanced green fluorescent protein (eGFP) trap in the gene’s first intron, *dilp8^MI00727^* (Garelli et al., 2012) (Fig. 1C). Tissue expression data indicates that in adults the *dilp8* transcript is high in females, where it is seen predominantly in ovaries, whereas male expression is overall very low [modENCODE_mRNA-Seq_tissues, (Brown et al., 2014); http://flyatlas.gla.ac.uk/FlyAtlas2/index.html, (Leader et al., 2017)]. Thus, we analyzed only females and observed *dilp8*-GFP in follicle cells in egg chambers of ovaries in 1-7 d old flies (Fig. 1A). DILP8 immunolabeling was detected in the same cell type (Fig. 1B). To control for antiserum-specificity we used *dilp8* mutant flies (*dilp8^MI00727^*) with a GFP insertion and could show that the DILP8 antiserum no longer recognized the GFP-labeled follicle cells (Fig. 1C, D). As another control for the specificity of the anti-DILP8 we show here that it labels punctates in wing imaginal discs after induction of disrupted development [see (Colombani et al., 2012;Garelli et al., 2012)] by knockdown of syntaxin 7 (*avalanche; avl*) through driving *avl*-RNAi with an *egrailed*-Gal4 driver (Supplementary figure 1).

**Fig. 1.**
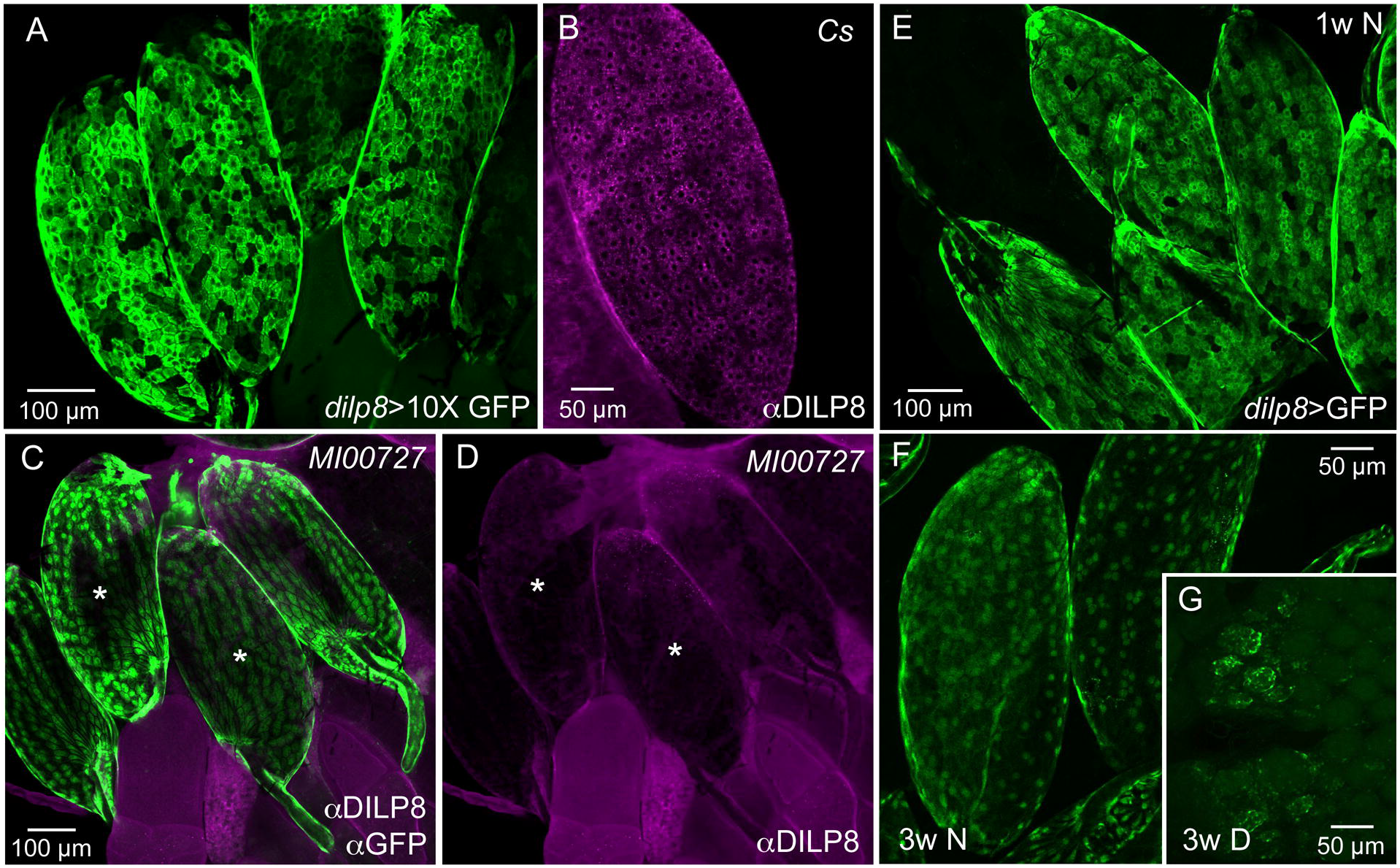
Ovarian follicle cells express *dilp8*/DILP8. **A.** Using a 10X-GFP the *dilp8* expression is strong in follicle cells surrounding mature eggs. **B.** DILP8 antiserum labels follicle cells in a wild type fly (*Cs, Canton s*). **C.** A *dilp8* mutant fly (*dilp8^MI00727^*) with a GFP-insertion reveals GFP expression in follicle cells. The sample was also incubated in anti-DILP8. **D.** DILP8 antiserum yields no immunolabeling in the same specimen due to the *dilp8* mutation. Asterisks label the same two eggs in C and D. **E.** In one-week-old flies kept under normal conditions *dilp8* expression is seen in follicle cells. **F.** In three-week-old flies *dilp8* expression is somewhat weaker. **G**. In flies kept for three weeks in reproductive diapause ovary development is arrested and only residual dilp8 expression is seen.

The *dilp8-*GFP expression in follicle cells is prominent in young flies, but can still be seen in three-week-old ones (Fig. 1E, F). In flies kept for three weeks in reproductive diapause, sparse *dilp8*-GFP is seen in cells of the rudimentary ovaries (Fig. 1G). A decrease in *dilp8* expression during adult diapause was shown earlier in a genome-wide analysis of gene transcripts (Kucerova et al., 2016).

We could not detect DILP8 immunolabeling outside the ovaries in adult flies under normal conditions. However, *dilp8*-expression could be seen in *dilp8*>GFP flies in cells of the hindgut and the salivary gland (Supplementary figure 2A, B). The gut expression is supported by single cell transcriptomics data where *dilp8* was found in intestinal muscle cells and enteroblasts [http://flygutseq.buchonlab.com/data; (Buchon et al., 2013)]. In third instar larvae, no DILP8 immunolabeling was seen unless imaginal discs were damaged (Supplementary figure 1), but *dilp8*-driven GFP was detected in neuronal progenitors in the brain (Supplementary figure 2C, D). Thus, under normal conditions weak DILP8 immunolabeling only appears in follicle cells and we additionally provide two lines of evidence for *dilp8* expression in these cells, dilp8>GFP and the eGFP insertion in the *dilp8^MI00727^* line. Recently, it was also shown by single-cell transcriptome sequencing that *dilp8* transcript displays a peak expression in late stage 14 follicle cells (Jevitt et al., 2020).

### Distribution of the DILP8 receptor Lgr3

The widespread distribution of the DILP8 receptor, Lgr3, has been described in the central nervous system (CNS) of larvae (Colombani et al., 2015;Garelli et al., 2015;Vallejo et al., 2015;Jaszczak et al., 2016) and adults (Meissner et al., 2016). These studies show numerous *Lgr3*-expressing neurons in the CNS. We used several *Lgr3*-Gal4 lines and could confirm the distribution in the brain and ventral nerve cord of adults (Fig. 2A-E). Some of the brain *Lgr3* neurons have processes in the pars intercerebralis (PI) and tritocerebrum/subesophageal zone (Fig. 2A-D) where also insulin producing cells (IPCs) and other median neurosecretory cells (MNCs) arborize [see (Cao and Brown, 2001;Nässel and Zandawala, 2019)]. To test whether the *Lgr3* expressing neurons are in synaptic contact with DILP2,3,5-producing IPCs we employed GFP-reconstitution across synaptic partners (GRASP) technique (Feinberg et al., 2008;Gordon and Scott, 2009). Using a R19B09-Gal4 and a dilp2-LexA line as well as UAS-spGFP1-10 and LexAop-spGFP11 we could show that in the larval brain IPCs are connected to *Lgr3*-expressing neurons (Supplementary figure 3A, B). However in the adult brain we found no reconstituted GFP (Supplementary figure 3C, D) suggesting the neuron types are no longer connected.

**Fig. 2.**
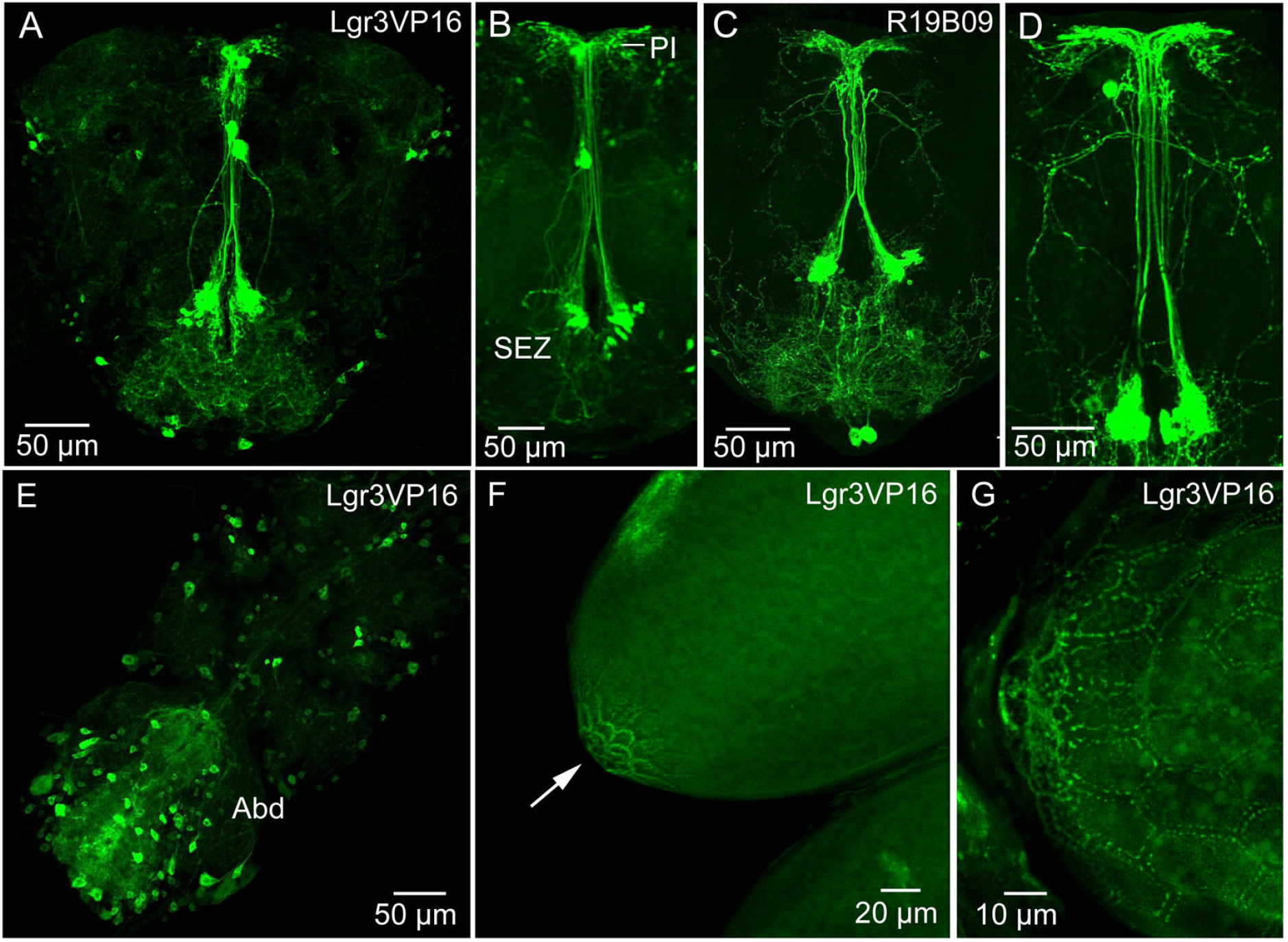
Expression of *Lgr3* in CNS and ovaries. **A**. Lgr3 expression in neurons of the adult brain using the Lgr3VP16-Gal4 driver. **B.** The same driver in a different brain showing *Lgr3* neurons of the subesophageal zone (SEZ) that supply processes to the pars intercerebralis. **C** and **D**. Another *Lgr3* driver (R19B09) displays a similar expression. **E.** Numerous neurons in the adult ventral nerve cord express *Lgr3*, especially in the abdominal neuromeres (Abd). **F** and **G**. *Lgr3* expression in follicle cells in the area closest to the oviduct (arrow in F). A detail at higher magnification from another ovary is shown in G.

We examined Lgr3 expression also in the CNS of larvae (Supplementary figure 4A, B). A set of 14 *Lgr3*-expressing neurons in the larval abdominal neuromeres co-express the neuropeptide leucokinin (LK) (Supplementary figure 4A, B). In adults, these neurons have been shown to use LK to regulate secretion in Malpighian tubules in control of ion and water homeostasis [(Zandawala et al., 2017); see also (Terhzaz et al., 1999)]. However, we could not detect colocalized Lgr3 and LK in these cells in adults. Importantly, we found *Lgr3*-Gal4 expression in a subset of follicle cells located closest to the oviduct (Fig. 2F, G). Furthermore, we detected *Lgr3* expression in a small set of peripheral bipolar neurons that innervate the dorsal aorta (heart) in the abdomen (Supplementary figure 4C-H). A median pair of such *Lgr3*-expressing aorta-associated cells co-express ion transport peptide (ITP) (Supplementary figure 4E-H). Similar median cells were identified by ITP antiserum several years ago (Dircksen et al., 2008).

Looking at the *Lgr3*-expressing neurons of the abdominal neuromeres in more detail we detected axons running towards the periphery (Fig. 3A-D). Since it is known that the ovaries and oviduct are innervated by octopaminergic neurons (Monastirioti, 2003;Rubinstein and Wolfner, 2013;Pauls et al., 2018), we performed double labeling with *Lgr3*-Gal4-driven GFP and immunostaining with antiserum to TDC2, a biosynthetic enzyme catalyzing the production of octopamine (OA) and tyrosine (Tyr). We detected TDC2-immunlabeled axons running posteriorly from abdominal neuromeres (Fig. 3C, D) and a few of these axons co-express *Lgr3* and TDC2 (Fig 3D). However, when analyzing the expression in the ovaries and oviduct, we found separate *Lgr3* and TDC2 expressing axons and no colocalization (Fig. 3E, F). Thus, we identified some *Lgr3*-expressing axons innervating oviduct muscle, but have no evidence that these *Lgr3* neurons also produce OA or Tyr.

**Fig. 3.**
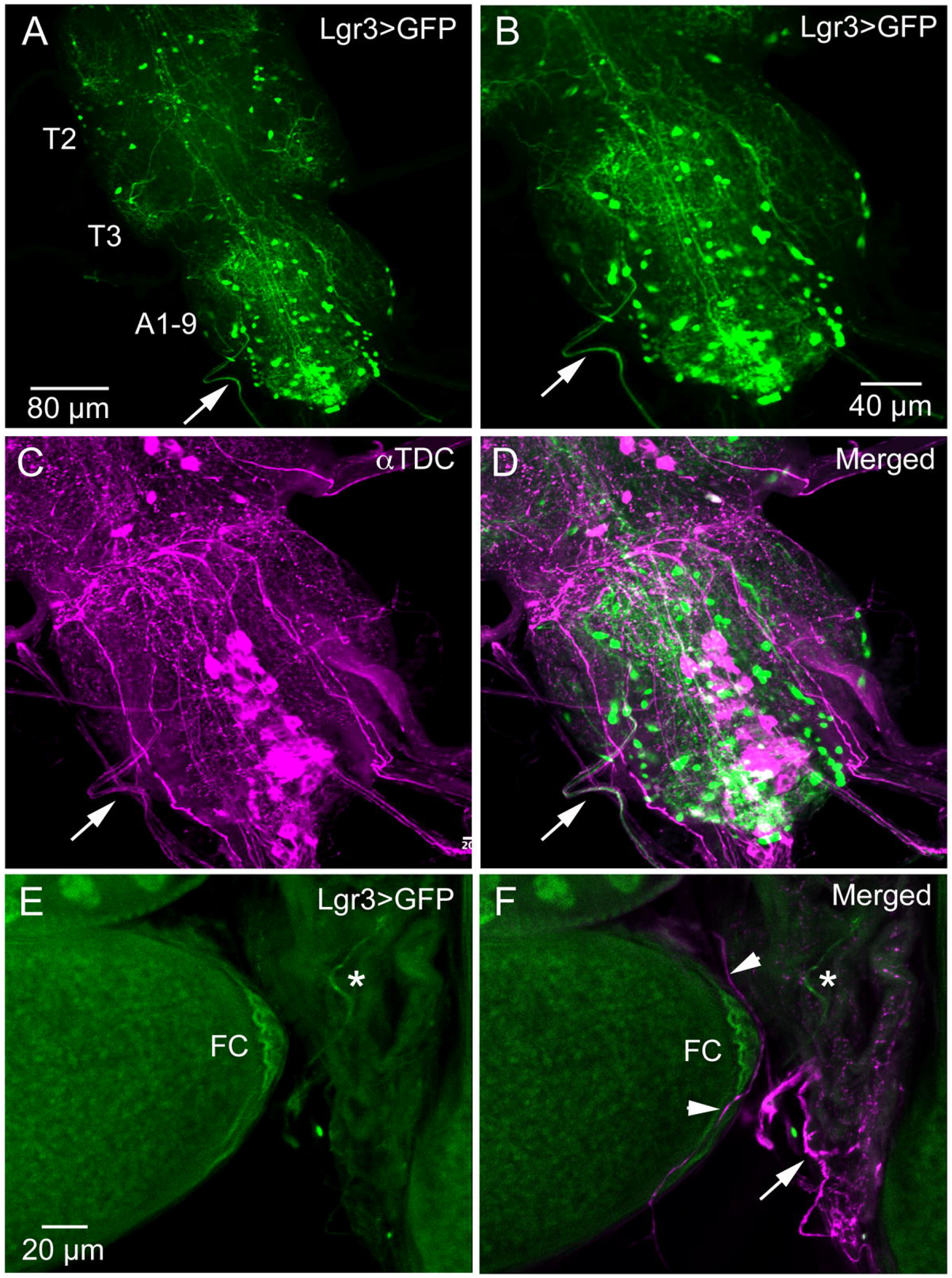
*Lgr3* and octopamine expressing efferent neurons innervate oviduct muscle. **A.** Overview of adult thoracic (T3, T3) and abdominal (A1-9) neuromeres with Lgr3 expressing neurons. One efferent axon emerging from the abdominal neuromeres is seen at arrow (Lgr3VP16-Gal4). **B-D.** The same specimen, which has also been immunolabeled with antiserum to Tdc2 (tyrosine decarboxylase; magenta). Note that the peripheral axon at arrows colocalizes Tdc2 and *Lgr3* labeling. Also note that quite a few Tdc2 immunolabeled peripheral axons emerge from the ventral nerve cord, some destined for the oviduct muscle. **E** and **F**. Part of the ovary and oviduct muscle with Lgr3 and Tdc2 expression. Basal follicle cells (FC) express *Lgr3* and so does one axon on oviduct muscle (at asterisk). In F several axons immunolabeled with anti-Tdc2 are seen on oviduct muscle (arrow and arrow heads). No colocalization of Lgr3 and Tdc2 could be detected.

### Functional roles of DILP8 signaling

Next, we addressed the functional roles of DILP8 signaling in adults. Since several of the DILPs are known to affect stress responses (Broughton et al., 2005;Grönke et al., 2010;Owusu-Ansah and Perrimon, 2015;Nässel and Vanden Broeck, 2016), we explored the role of *dilp8* in this respect. Using the *dilp8* mutant flies (*dilp8^MI00727^*), we tested the effect loss of *dilp8* on resistance to starvation (Fig. 4A) and desiccation (Fig. 4B). In both tests the mutant flies survived longer than control flies. To analyze the effect of *dilp8* gain of function we used different Gal4 lines to drive UAS-*dilp8*. The efficacy of these lines is shown in Supplementary figure 5. We used a Gal4 line *c929* that drives expression in several hundred peptidergic neurons characterized by the transcription factor *Dimmed* (Hewes et al., 2003). With the *c929*-Gal4 a substantial number of *Dimmed* neurons could be labeled with antiserum to DILP8 (Supplementary figure 5A, B). It can be noted that *dilp8* overexpression leads to a down-regulation of DILP3 expression in insulin producing cells (IPCs) of the brain (Supplementary figure 5C-E) and possibly affects other DILPs (not tested). Also a *dilp2*-Gal4 is able to induce DILP8 immunolabeling in IPCs (Supplementary figure 5F).

**Fig. 4.**
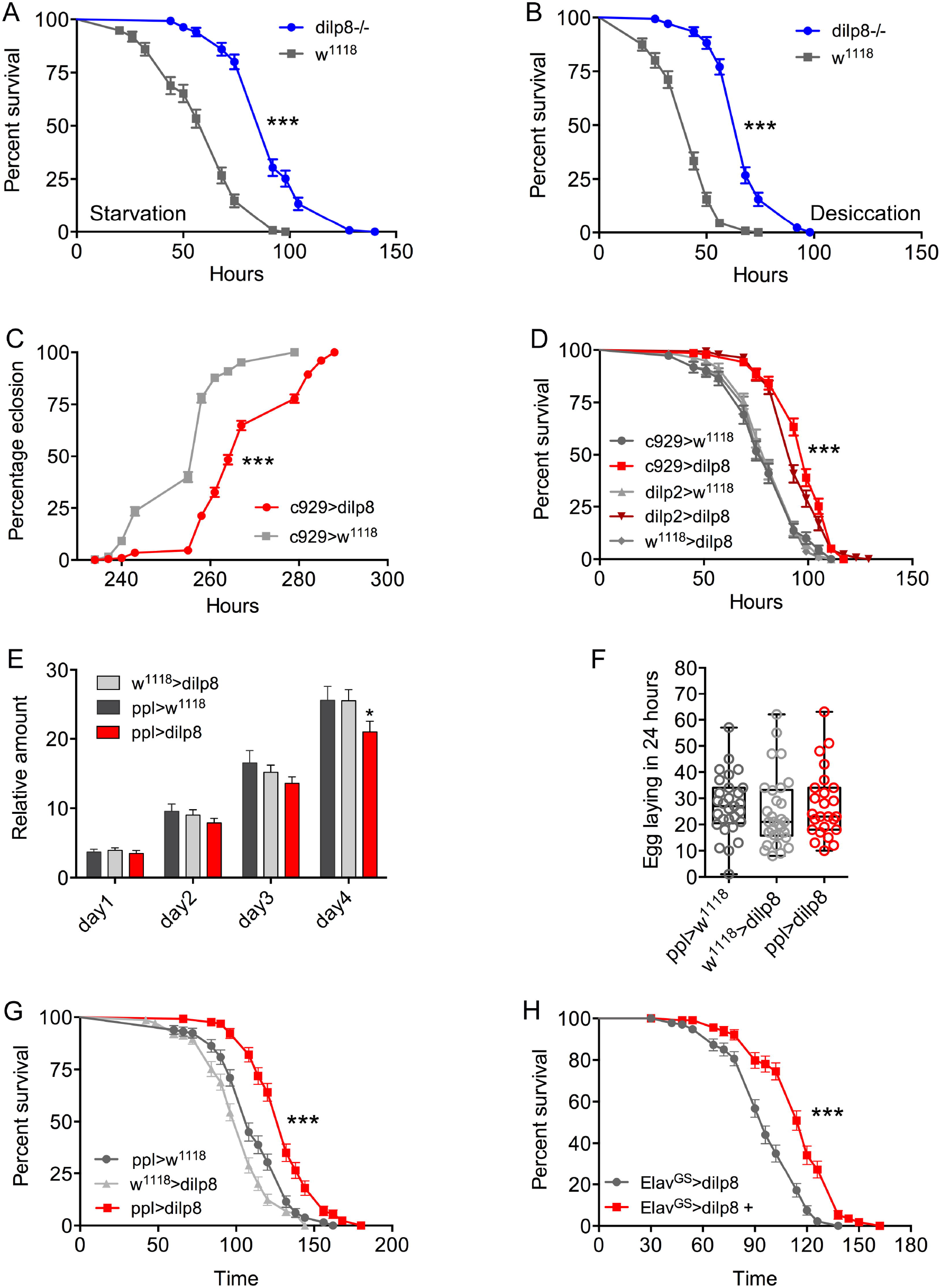
Manipulations of *dilp8* affect stress responses, feeding and body weight. **A** and **B**. The *dilp8* mutant flies survive starvation and desiccation better than controls. We used 135 flies per genotype from three independent replicates [***p<0.001, as assessed by log-rank (Mantel–Cox) test]. **C.** Overexpression of *dilp8* with the *c929*-Gal4 driver delays adult eclosion (time from egg to eclosion). The eclosion time of more than 394 flies were analyzed here, [***p<0.001, as assessed by log-rank (Mantel–Cox) test]. **D.** Overexpression of *dilp8* with the *c929-* and *dilp2*-Gal4 lines increase resistance to desiccation. More than 110 flies from three independent replicates were used [***p<0.001, as assessed by log-rank (Mantel–Cox) test]. **E**. Ectopic expression of *dilp8* in the fat body (*ppl*-Gal4) decreases food intake as monitored by CAFE assay. Cumulative data for four days are shown. More than 96 flies from three independent replicates were used [***p<0.001, as assessed by log-rank (Mantel–Cox) test]. **F.** Egg laying over 24 h is not affected by *dilp8* expression in the fat body. 25-30 flies from 3 replicates were used (one-way ANOVA followed by Tukey’s test). **G**. Ectopic expression of *dilp8* in the fat body increases resistance to starvation. More than 96 flies from three independent replicates were used [***p<0.001, as assessed by log-rank (Mantel–Cox) test]. **H.** Using a drug-inducible panneuronal driver, gene-switch *Elav*-Gal4 (*Elav^GS^*-Gal4) we could induce *dilp8* in five-day-old flies by feeding RU486 (Elav^GS^-Gal4>dilp8+). Ectopic *dilp8* increases starvation resistance. More than 96 flies from three independent replicates were used [***p < 0.001, as assessed by log-rank (Mantel–Cox) test].

As seen in Figure 4C, over-expression of *dilp8* with the *c929*-Gal4 delays adult eclosion (time from egg to adult eclosion increases). This is as expected from the role of *dilp8* in inhibiting Ecd production and thereby causing developmental delay (Colombani et al., 2012;Garelli et al., 2012). Both *c929*- and *dilp2*-Gal4 driven *dilp8* results in increased survival during starvation (Fig. 4D). We next employed a driver, *pumpless* (*ppl*) that directs expression to the fat body. Using *ppl>dilp8* we noted that one-week-old flies were feeding less over four days (Fig. 4E). *The ppl>dilp8* flies did not display any difference in egg laying (Fig. 4F), but survived starvation better than controls (Fig. 4G). To test the temporally restricted effect of *dilp8* overexpression in adult flies, we employed *Elav^GS^*-Gal4 flies where the Gal4 is activated by feeding the flies RU486 and found that flies display increased starvation resistance (Fig. 4H; Supplementary figure 5G, H). We furthermore tested broad ectopic *dilp8* expression driven with a *daughterless* (*Da*) gene-switch Gal4 and obtained a similar adult-specific result (Supplementary figure 5I-L). These conditional experiments confirm that broad ectopic *dilp8* expression increases starvation resistance in adults.

Inspired by the expression of *dilp8*/DILP8 in follicle cells and *Lgr3* in a subset of follicle cells in the basal part of the egg chamber, as well as in axons innervating the oviduct, we went on to test the role of *dilp8* in fecundity. The number of eggs laid is significantly diminished in *dilp8* mutant flies (Fig. 5A). This reduced number appears to be a result of defects in ovulation since the number of eggs retained in the ovaries is higher in *dilp8* mutant flies (Fig. 5B, C). As seen in Figure 5B, the increased number of eggs in ovaries is reflected in an enlarged abdomen in the mutant flies. We also overexpressed and knocked down *dilp8* with a *dilp8*-Gal4 driver, but only noted a significant reduction of eggs laid for *dilp8*>*dilp8*-RNAi (Fig. 5D). Next, we tested the effects of manipulating *dilp8* levels specifically in follicle cells by overexpression or knockdown with the *FC2*-Gal4 driver (Fig. 5E, F). These manipulations did not significantly affect the number of eggs laid in 48 h, but the number of eggs remaining in the ovaries increased after *dilp8*-RNAi (Fig. 5F). Thus, *dilp8* in follicle cells appears critical for ovulation.

**Fig. 5.**
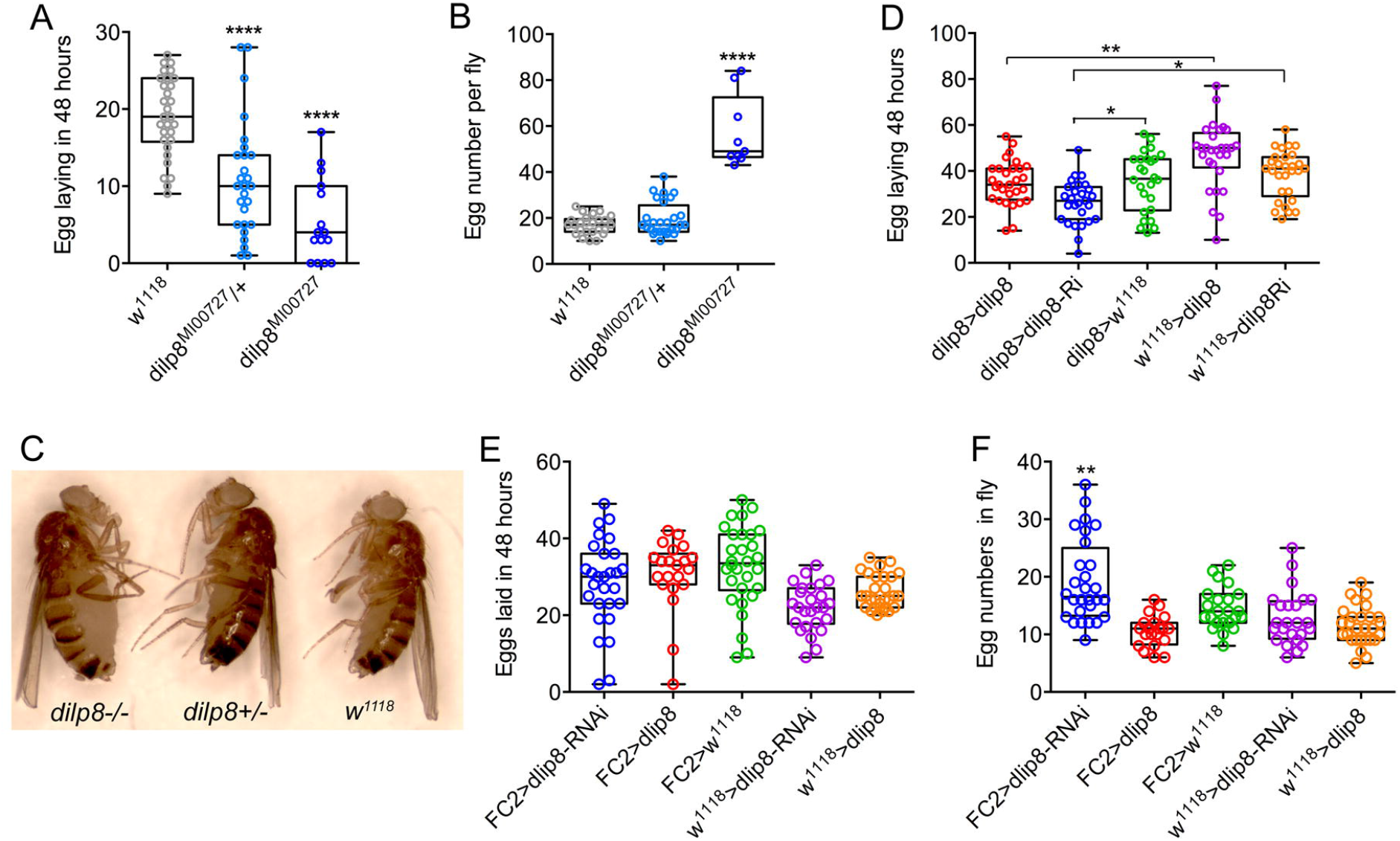
Manipulations of *dilp8* affect fecundity in flies. **A.***dilp8* mutants (hetero- and homozygous) lay fewer eggs than controls. 15-30 flies from 3 replicates were used (***p < 0.001, one-way ANOVA followed by Tukey’s test). **B.** The number of eggs retained in the flies is significantly higher in homozygous *dilp8* mutants. 9-27 flies from 3 replicates were used (***p < 0.001, one-way ANOVA followed by Tukey’s test). **C**. The higher number of eggs retained in *dilp8* mutant flies is reflected in the bloated abdomen (not seen in heterozygote or control). **D**. Knockdown of *dilp8* using *dilp8*-Gal4 significantly reduces egg laying over 48 h, whereas *dilp8* overexpression has no effect. E. Overexpression and knockdown of dilp8 in follicle cells (*FC2*-Gal4) does not significantly affect egg laying over 48 h. 21-30 flies from 3 replicates were used (one-way ANOVA followed by Tukey’s test). **F.** Knockdown of dilp8 in follicle cells leads to a higher number of eggs retained in flies, but overexpression has no effect [20-28 flies from 3 replicates were used (***p < 0.001, one-way ANOVA followed by Tukey’s test)].

## Discussion

Our study suggests that *dilp8* in follicle cells of ovaries is important for fecundity in *Drosophila*. We could show *dilp8*-Gal4 expression and DILP8 immunolabeling in follicle cells of young mated and unmated flies. Flies mutant in *dilp8* lay fewer eggs, but seem to produce normal numbers of eggs that are retained within the ovaries, suggesting effects on ovulation. Knockdown of *dilp8* specifically in follicle cells also resulted in flies with more eggs retained in the ovaries, (but we did not see fewer eggs laid). Since we could not detect both *dilp8* and DILP8 expression outside the ovaries in adult flies under normal conditions, we suggest that follicle cells are a primary source of the peptide hormone in regulation of fecundity. The target of DILP8 appears to be *Lgr3*-expressing neurons in the brain and ventral nerve cord, as well as a subset of the follicle cells found closest to the oviduct (see Fig. 6). We did find some *Lgr3*-expressing efferent neurons in the abdominal neuromeres that send axons to the oviduct muscle. Thus DILP8 could act both in a paracrine fashion directly on follicle cells to induce rupture and subsequent ovulation and act systemically on neurons innervating the oviduct muscle that regulate ovulation. Since octopaminergic neurons innervating the oviduct have been implicated in regulation of ovulation (Monastirioti, 2003;Lee et al., 2009;Rubinstein and Wolfner, 2013;Sun and Spradling, 2013;Deady et al., 2015;Pauls et al., 2018), and follicle cell maturation [see (Knapp et al., 2020)], we tested whether octopaminergic neurons express the *Lgr3* receptor. Indeed a few abdominal efferents were found to coexpress *Lgr3* and immunolabeling for TDC2, the biosynthetic enzyme of octopamine and tyramine. However, these did not seem to innervate oviduct muscle, so it does not appear that DILP8 acts directly on octopaminergic neurons that regulate ovulation. The neurotransmitter of the efferent *Lgr3* neurons that supply oviduct muscle, thus, remains to be identified.

**Fig. 6.**
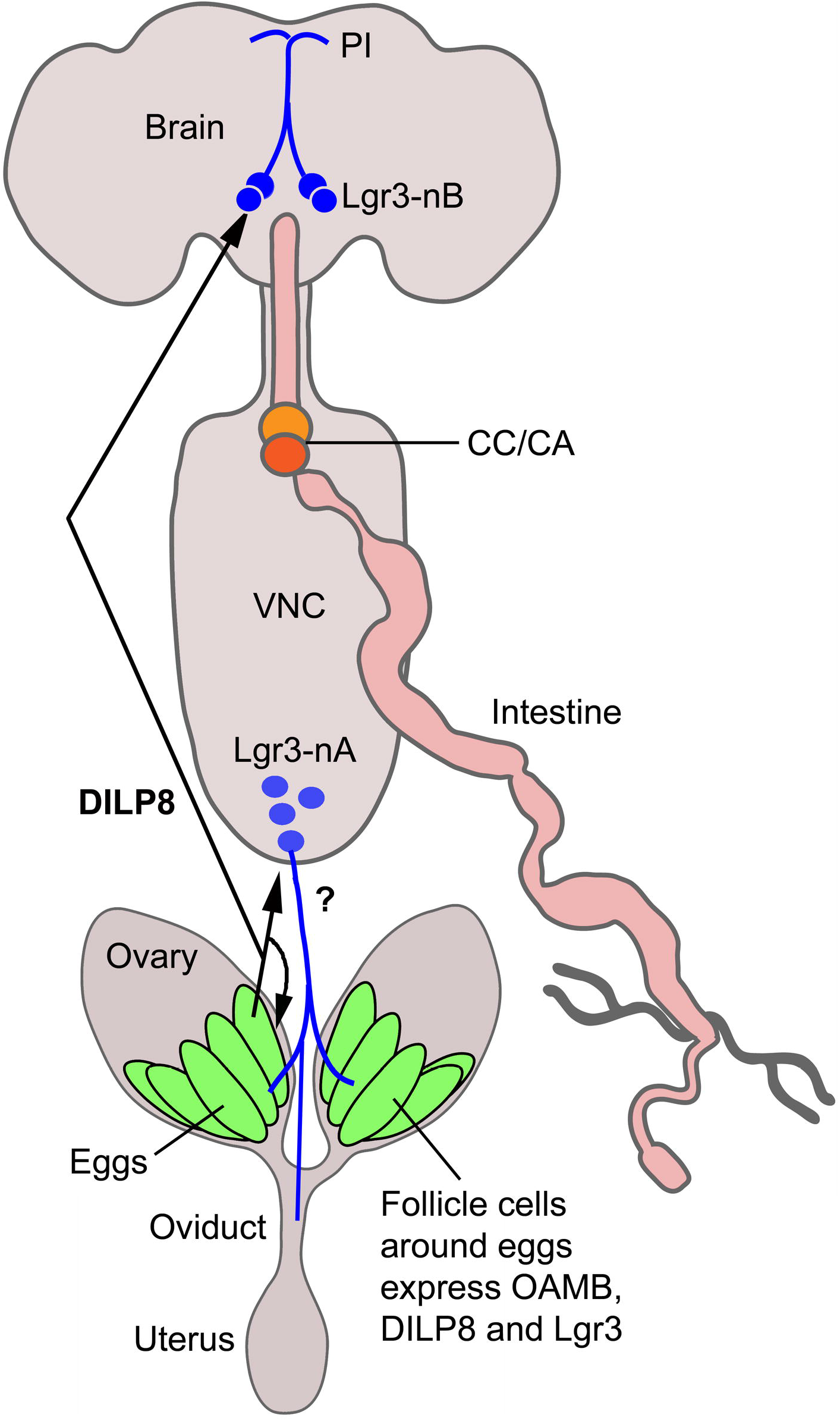
Summary diagram of DILP8 signaling in adult *Drosophila*. DILP8 is produced in follicle cells of the ovaries and can act on the receptor Lgr3 expressed on neurons of the CNS (blue) and on follicle cells near the oviduct. The efferent neurons Lgr3-nA in the ventral nerve cord (VNC) innervate the oviduct, but we found no evidence for octopamine (OA) coexpression, so the neurotransmitter is unknown (?). Other efferent neurons have been shown to produce OA and regulate ovulation via the OA receptor OAMB. A set of Lgr3 neurons (Lgr3-nB) in the brain supplies processes to the pars intercerebralis (PI) where IPCs and other median neurosecretory cells (MNCs) have dendrites. It is tempting to speculate that DILP8 acts on such neurons and that these in turn regulate release of DILPs or other peptide hormones in MNCs to regulate feeding and metabolism. CC/CA, corpora cardiaca/corpora allata.

It is remarkable that *dilp8*/DILP8 is restricted to follicle cells, as also indicated by available RNA expression data [FlyAtlas2 and ModEncode (Brown et al., 2014;Leader et al., 2017)]. We do see *dilp8*-GFP expression also in cells of the hindgut and salivary glands, but could not find DILP8-immunolabeling in those tissues. Thus, adult expression is quite restricted, as is the case in larvae where DILP8 can be induced in imaginal discs, but not elsewhere in the organism it seems (Colombani et al., 2012;Garelli et al., 2012) (see also Supplementary figure 1). In contrast, the DILP8 receptor *Lgr3* is expressed in a relatively large number of neurons in the brain and ventral nerve cord, both in larvae and adults (Colombani et al., 2015;Garelli et al., 2015;Vallejo et al., 2015;Meissner et al., 2016). So far only a specific set of *Lgr3* cells in the larval brain have been explored functionally. These two pairs of cells [growth coordinating Lgr3 (GCL) neurons] innervate the four neurons producing PTTH and thereby regulate ecdysone production in the prothoracic gland to produce a developmental delay (Colombani et al., 2015;Garelli et al., 2015;Vallejo et al., 2015). In adult flies, activation of sex-specific *Lgr3* neurons in the abdominal neuromeres inhibits female reproductive receptivity and fecundity (Meissner et al., 2016). We did not analyze effects of *Lgr3* manipulations here, but suggest that this receptor in follicle cells and perhaps in a small set of efferent *Lgr3* neurons innervating oviduct muscle may be primarily responsible for the ovulation phenotype. It cannot be excluded that other *Lgr3*-expressing neurons in the abdominal ganglia interact with octopaminergic neurons innervating the oviduct and thus indirectly contribute to the known octopamine effects on ovary maturation and ovulation (Lee et al., 2003;Monastirioti, 2003;Lee et al., 2009;Rubinstein and Wolfner, 2013;Deady et al., 2015;Yoshinari et al., 2020).

We noted effects of *dilp8* manipulations on resistance to starvation and food intake, which might suggest that the follicle cells signal systemically to regulate metabolism. Possibly this reflects feedback nutritional signaling from the ovaries to brain neurosecretory cells to ensure allocation of nutrients. There is no *Lgr3* expression in brain insulin-producing cells (IPCs) or other MNCs, and in adults we failed to demonstrate direct connections between *Lgr3* neurons and IPCs using the GRASP technique. However, in larvae, a connection between these neurons was seen. There are however, sets of *Lgr3* neurons with processes in pars intercerebralis where IPCs have dendrites (see Fig. 2A-D). Thus, DILP8 might act on these *Lgr3* neurons and they may in turn signal to IPCs or other MNCs in a paracrine (non synaptic) fashion. Alternatively DILP8 might act directly on IPCs or other MNCs via the tyrosine kinase insulin receptor, dInR. It was shown that dInR could be captured by DILP8 in a ligand capture assay, possibly suggesting that DILP8 could bind to both Lgr3 and dInR (Garelli et al., 2015).

Furthermore, we showed that *dilp8* mutants display increased resistance to desiccation, suggesting effects of DILP8 on water homeostasis. This could be mediated indirectly via action on *Lgr3*-expressing neurons in the abdominal ganglia that produce diuretic or antidiuretic hormones. In larvae we detected *Lgr3* expression in leucokinin-producing neurosecretory cells (ABLKs) in abdominal ganglia, known to regulate water homeostasis in adult flies (Zandawala et al., 2017). These neurons also express the dInR both in larvae and adults (Liu et al., 2015). However, in adults the *Lgr3*-leucokinin coexpression was not seen, but we found *Lgr3*-expession in a small set of peripheral neurons that produce ion transport peptide, an antidiuretic hormone (Dircksen et al., 2008;Galikova et al., 2018). The functional role of these neurons in water homeostasis remains to be demonstrated and it would be interesting to determine whether DILP8 regulates their activity.

In summary, we found that *dilp8*/DILP8 is expressed in follicle cells of the ovaries, and that fecundity and metabolism is affected by manipulations of *dilp8*. We propose that DILP8 signaling might represent a feedback from ovaries to the brain to ensure allocation of nutrients for egg maturation and that DILP8 also constitutes a signal that aids in ovulation in concert with octopamine. Thus, DILP8 appears to act as a relaxin-like hormone in regulation of fecundity, suggesting conservation of an ancient function from insects to mammals.

## Supplementary material figures

**Supplementary figure 1**. DILP8 immunoreactivity can be induced in wing imaginal discs by disrupting development. **A-C**. Using an *engrailed* (en) Gal4-driver (with GFP insertion) to knock down avalanche (en-GFP>avl-RNAi) we could induce DILP8 immunolabeling (framed area in B). GFP marks engrailed expression. **D-F.** Higher magnification of the area framed in B. **G-I.** Controls (en-GFP>w^1118^) where on DILP8 immunolabeling is seen (**H**).

**Supplementary figure 2**. Further *dilp8* expression in tissues of adult and larval *Drosophila*. **A.** Using dilp8-Gal4 to drive 10X-GFP we could demonstrate expression in follicle cells of ovary and in cells of hindgut. **C.** Expression of *dilp8* in cells of salivary gland of adult fly. **C** and **D**. In third instar larvae neuronal progenitor cells of the brain express *dilp8*. In **D** glial cells are shown with repo-immunolabeling. VNC, ventral nerve cord.

**Supplementary figure 3**. GFP-reconstitution across synaptic partners (GRASP) reveals contacts between Lgr3 neurons and IPCs. **A** and **B**. In the larval brain *dilp2*-LexA and R19B09-Gal4 generated GRASP (R19B09 Gal4>UAS-spGFP1-10; dilp2-lexA>LexAop-spGFP11; see methods) results in reconstituted GFP suggesting connections between IPCs and *Lgr3* expressing neurons (**B**). In A (and C) the red is red fluorescent protein (RFP) driven by *dilp2*-LexA; blue is DAPI labeling of nuclei. **C** and **D**. In adults GRASP with the same fly lines does not result in GFP, suggesting IPCs and Lgr3 neurons no longer form connections. Arrows in A and C points at cell bodies of IPCs (weakly labeled).

**Supplementary figure 4**. Lgr3 expression in larval CNS and adult peripheral neurons. **A** and **B**. In the larval abdominal neuromeres *Lgr3* is expressed in 7 pairs of leucokinin (LK)-producing neurosecretory cells (ABLKs) indicated by arrows. **C-H**. *Lgr3* is expressed in a small set of peripheral neurons. Some of these are lateral and segmental (s) and one pair is median (m), indicated by arrows in **C-E**). Both types of neurons innervate the dorsal aorta, or heart (see **D**). The segmental axons run along the alary muscles (am). The median *Lgr3* neurons express ion transport peptide (ITP) immunoreactivity (**F-H**). Note that the cell bodies are not clearly labeled with the ITP antiserum (**G** and **H**).

**Supplementary figure 5**. Overexpression of *dilp8* in different neurons results in DILP8 immunolabeling. **A**. The *c929*-Gal4>*dilp8* results in DILP8 immunolabeling in many peptidergic neurons, including median neurosecretory cells (MNC; see also enlarged at * in inset) and large neurons in the subesophageal zone (SEZ). **B.** In controls (c929>w^1118^) no DILP8 immunolabeling can be detected. **C** and **D**. *c929*-Gal4>*dilp8* results in decreased DILP3 immunolabeling. **E**. Quantification of DILP3 labeling [5 flies used for *c929*>*dilp8* and 10 flies used for *c929*>*w^1118^* (*p < 0.05, unpaired Student’s t-test)]. **F**. Ectopic expression of dilp8 in IPCs (dilp2>dilp8) results in DILP8 immunoreactivity. **G** and **H**. Conditional gene switch Gal4 expression (ElavGS>dilp8) induced in adult flies by feeding RU486 triggered DILP8 immunoreactivity in numerous neurons (**G**), and no labeling was seen without feeding RU486 (**H**). **I-K**. DILP8 immunolabeling in brain of *Da^GS^*>*dilp8* flies fed RU486 as adults. DILP8 immunolabeling can be seen in median neurosecretory cells (MNC), neurons of subesophageal zone (SEZ) and optic lobe (OL). **L.** Starvation resistance of *Da^GS^*>*dilp8* flies with RU486 activation (+) at adult stage increases compared to inactivated flies. 45 flies from three independent replicates were used [**p < 0.01, as assessed by log-rank (Mantel–Cox) test].

## References

Ahmad, M., He, L., and Perrimon, N. (2019). Regulation of insulin and adipokinetic hormone/glucagon production in flies. WIREs Developmental Biology, e360.

Brand, A.H., and Perrimon, N. (1993). Targeted gene expression as a means of altering cell fates and generating dominant phenotypes. Development 118, 401–415.

Brogiolo, W., Stocker, H., Ikeya, T., Rintelen, F., Fernandez, R., and Hafen, E. (2001). An evolutionarily conserved function of the *Drosophila* insulin receptor and insulin-like peptides in growth control. Curr Biol 11, 213–221.

Broughton, S.J., Piper, M.D., Ikeya, T., Bass, T.M., Jacobson, J., Driege, Y., Martinez, P., Hafen, E., Withers, D.J., Leevers, S.J., and Partridge, L. (2005). Longer lifespan, altered metabolism, and stress resistance in *Drosophila* from ablation of cells making insulin-like ligands. Proc Natl Acad Sci U S A 102, 3105–3110.

Brown, J.B., Boley, N., Eisman, R., May, G.E., Stoiber, M.H., Duff, M.O., Booth, B.W., Wen, J., Park, S., Suzuki, A.M., Wan, K.H., Yu, C., Zhang, D., Carlson, J.W., Cherbas, L., Eads, B.D., Miller, D., Mockaitis, K., Roberts, J., Davis, C.A., Frise, E., Hammonds, A.S., Olson, S., Shenker, S., Sturgill, D., Samsonova, A.A., Weiszmann, R., Robinson, G., Hernandez, J., Andrews, J., Bickel, P.J., Carninci, P., Cherbas, P., Gingeras, T.R., Hoskins, R.A., Kaufman, T.C., Lai, E.C., Oliver, B., Perrimon, N., Graveley, B.R., and Celniker, S.E. (2014). Diversity and dynamics of the Drosophila transcriptome. Nature 512, 393.

Buchon, N., Osman, D., David, F.P., Fang, H.Y., Boquete, J.P., Deplancke, B., and Lemaitre, B. (2013). Morphological and molecular characterization of adult midgut compartmentalization in *Drosophila*. Cell Rep 3, 1725–1738.

Cao, C., and Brown, M.R. (2001). Localization of an insulin-like peptide in brains of two flies. Cell Tissue Res 304, 317–321.

Chintapalli, V.R., Wang, J., and Dow, J.A. (2007). Using FlyAtlas to identify better *Drosophila melanogaster* models of human disease. Nat Genet 39, 715–720.

Colombani, J., Andersen, D.S., Boulan, L., Boone, E., Romero, N., Virolle, V., Texada, M., and Leopold, P. (2015). Drosophila Lgr3 Couples Organ Growth with Maturation and Ensures Developmental Stability. Current biology: CB 25, 2723–2729.

Colombani, J., Andersen, D.S., and Leopold, P. (2012). Secreted peptide Dilp8 coordinates *Drosophila* tissue growth with developmental timing. Science 336, 582–585.

Deady, L.D., Shen, W., Mosure, S.A., Spradling, A.C., and Sun, J. (2015). Matrix metalloproteinase 2 is required for ovulation and corpus luteum formation in *Drosophila*. PLoS Genet 11, e1004989.

Dircksen, H., Tesfai, L.K., Albus, C., and Nässel, D.R. (2008). Ion transport peptide splice forms in central and peripheral neurons throughout postembryogenesis of *Drosophila melanogaster*. J Comp Neurol 509, 23–41.

Feinberg, E.H., Vanhoven, M.K., Bendesky, A., Wang, G., Fetter, R.D., Shen, K., and Bargmann, C.I. (2008). GFP Reconstitution Across Synaptic Partners (GRASP) Defines Cell Contacts and Synapses in Living Nervous Systems. Neuron 57, 353–363.

Galikova, M., Dircksen, H., and Nässel, D.R. (2018). The thirsty fly: Ion transport peptide (ITP) is a novel endocrine regulator of water homeostasis in *Drosophila*. PLoS Genet 14, e1007618.

Garelli, A., Gontijo, A.M., Miguela, V., Caparros, E., and Dominguez, M. (2012). Imaginal discs secrete insulin-like peptide 8 to mediate plasticity of growth and maturation. Science 336, 579–582.

Garelli, A., Heredia, F., Casimiro, A.P., Macedo, A., Nunes, C., Garcez, M., Dias, A.R., Volonte, Y.A., Uhlmann, T., Caparros, E., Koyama, T., and Gontijo, A.M. (2015). Dilp8 requires the neuronal relaxin receptor Lgr3 to couple growth to developmental timing. Nat Commun 6, 8732.

Gontijo, A.M., and Garelli, A. (2018). The biology and evolution of the Dilp8-Lgr3 pathway: A relaxin-like pathway coupling tissue growth and developmental timing control. Mechanisms of Development 154, 44–50.

Gordon, M.D., and Scott, K. (2009). Motor control in a Drosophila taste circuit. Neuron 61, 373–384.

Grönke, S., Clarke, D.F., Broughton, S., Andrews, T.D., and Partridge, L. (2010). Molecular evolution and functional characterization of *Drosophila* insulin-like peptides. PLoS Genet 6, e1000857.

Hewes, R.S., Park, D., Gauthier, S.A., Schaefer, A.M., and Taghert, P.H. (2003). The bHLH protein Dimmed controls neuroendocrine cell differentiation in *Drosophila*. Development 130, 1771–1781.

Ja, W.W., Carvalho, G.B., Mak, E.M., De La Rosa, N.N., Fang, A.Y., Liong, J.C., Brummel, T., and Benzer, S. (2007). Prandiology of *Drosophila* and the CAFE assay. Proc Natl Acad Sci USA 104, 8253–8256.

Jaszczak, J.S., Wolpe, J.B., Bhandari, R., Jaszczak, R.G., and Halme, A. (2016). Growth Coordination During *Drosophila melanogaster* Imaginal Disc Regeneration Is Mediated by Signaling Through the Relaxin Receptor Lgr3 in the Prothoracic Gland. Genetics 204, 703.

Jevitt, A., Chatterjee, D., Xie, G., Wang, X.-F., Otwell, T., Huang, Y.-C., and Deng, W.-M. (2020). A single-cell atlas of adult *Drosophila* ovary identifies transcriptional programs and somatic cell lineage regulating oogenesis. PLOS Biology 18, e3000538.

Juarez-Carreño, S., Morante, J., and Dominguez, M. (2018). Systemic signalling and local effectors in developmental stability, body symmetry, and size. Cell stress 2, 340–361.

Knapp, E.M., Li, W., Singh, V., and Sun, J. (2020). Nuclear receptor Ftz-f1 promotes follicle maturation and ovulation partly via bHLH/PAS transcription factor Sim. eLife 9, e54568.

Kubrak, O.I., Kucerova, L., Theopold, U., and Nässel, D.R. (2014). The Sleeping Beauty: How Reproductive Diapause Affects Hormone Signaling, Metabolism, Immune Response and Somatic Maintenance in *Drosophila melanogaster*. PLoS ONE 9, e113051.

Kucerova, L., Kubrak, O.I., Bengtsson, J.M., Strnad, H., Nylin, S., Theopold, U., and Nässel, D.R. (2016). Slowed aging during reproductive dormancy is reflected in genome-wide transcriptome changes in *Drosophila melanogaster*. BMC genomics 17, 50.

Leader, D.P., Krause, S.A., Pandit, A., Davies, S.A., and Dow, J.A t. (2017). FlyAtlas 2: a new version of the Drosophila melanogaster expression atlas with RNA-Seq, miRNA-Seq and sex-specific data. Nucleic Acids Research 46, D809–D815.

Lee, H.G., Rohila, S., and Han, K.A. (2009). The octopamine receptor OAMB mediates ovulation via Ca2+/calmodulin-dependent protein kinase II in the Drosophila oviduct epithelium. PLoS One 4, e4716.

Lee, H.G., Seong, C.S., Kim, Y.C., Davis, R.L., and Han, K.A. (2003). Octopamine receptor OAMB is required for ovulation in *Drosophila melanogaster*. Dev Biol 264, 179–190.

Li, Q., and Gong, Z. (2015). Cold-sensing regulates *Drosophila* growth through insulin-producing cells. Nat Commun 6, 10083.

Liu, Y., Liao, S., Veenstra, J.A., and Nässel, D.R. (2016). *Drosophila* insulin-like peptide 1 (DILP1) is transiently expressed during non-feeding stages and reproductive dormancy. Sci Rep 6, 26620.

Liu, Y., Luo, J., Carlsson, M.A., and Nässel, D.R. (2015). Serotonin and insulin-like peptides modulate leucokinin-producing neurons that affect feeding and water homeostasis in *Drosophila*. J Comp Neurol 523, 1840–1863.

Meissner, G.W., Luo, S.D., Dias, B.G., Texada, M.J., and Baker, B.S. (2016). Sex-specific regulation of *Lgr3* in *Drosophila* neurons. Proceedings of the National Academy of Sciences 113, E1256.

Monastirioti, M. (2003). Distinct octopamine cell population residing in the CNS abdominal ganglion controls ovulation in *Drosophila melanogaster*. Dev Biol 264, 38–49.

Nässel, D.R., Cantera, R., and Karlsson, A. (1992). Neurons in the cockroach nervous system reacting with antisera to the neuropeptide leucokinin I. J Comp Neurol 322, 45–67.

Nässel, D.R., and Vanden Broeck, J. (2016). Insulin/IGF signaling in Drosophila and other insects: factors that regulate production, release and post-release action of the insulin-like peptides. Cell Mol Life Sci 73, 271–290

Nässel, D.R., and Zandawala, M. (2019). Recent advances in neuropeptide signaling in Drosophila, from genes to physiology and behavior. Progr Neurobiol 179, 101607.

Okamoto, N., Yamanaka, N., Yagi, Y., Nishida, Y., Kataoka, H., O’connor, M.B., and Mizoguchi, A. (2009). A fat body-derived IGF-like peptide regulates postfeeding growth in *Drosophila*. Dev Cell 17, 885–891.

Owusu-Ansah, E., and Perrimon, N. (2014). Modeling metabolic homeostasis and nutrient sensing in *Drosophila*: implications for aging and metabolic diseases. Disease models & mechanisms 7, 343–350.

Owusu-Ansah, E., and Perrimon, N. (2015). Stress Signaling Between Organs in Metazoa. Annual review of cell and developmental biology 31, 497–522.

Pauls, D., Blechschmidt, C., Frantzmann, F., and El Jundi, B. (2018). A comprehensive anatomical map of the peripheral octopaminergic/tyraminergic system of Drosophila melanogaster. 8, 15314.

Pfeiffer, B.D., Jenett, A., Hammonds, A.S., Ngo, T.T., Misra, S., Murphy, C., Scully, A., Carlson, J.W., Wan, K.H., Laverty, T.R., Mungall, C., Svirskas, R., Kadonaga, J.T., Doe, C.Q., Eisen, M.B., Celniker, S.E., and Rubin, G.M. (2008). Tools for neuroanatomy and neurogenetics in *Drosophila*. Proc Natl Acad Sci U S A 105, 9715–9720.

Ray, M., and Lakhotia, S.C. (2019). Activated Ras/JNK driven Dilp8 in imaginal discs adversely affects organismal homeostasis during early pupal stage in Drosophila, a new checkpoint for development. Developmental Dynamics 248, 1211–1231.

Rubinstein, C.D., and Wolfner, M.F. (2013). *Drosophila* seminal protein ovulin mediates ovulation through female octopamine neuronal signaling. Proceedings of the National Academy of Sciences 110, 17420.

Rulifson, E.J., Kim, S.K., and Nusse, R. (2002). Ablation of insulin-producing neurons in flies: growth and diabetic phenotypes. Science 296, 1118–1120.

Saunders, D.S., Henrich, V.C., and Gilbert, L.I. (1989). Induction of diapause in *Drosophila melanogaster*: photoperiodic regulation and the impact of arrhythmic clock mutations on time measurement. Proc Natl Acad Sci USA 86, 3748–3752.

Shimada, Y., Burn, K.M., Niwa, R., and Cooley, L. (2011). Reversible response of protein localization and microtubule organization to nutrient stress during *Drosophila* early oogenesis. Dev Biol 355, 250–262.

Slaidina, M., Delanoue, R., Grönke, S., Partridge, L., and Leopold, P. (2009). A *Drosophila* insulin-like peptide promotes growth during nonfeeding states. Dev Cell 17, 874–884.

Sun, J., and Spradling, A.C. (2013). Ovulation in Drosophila is controlled by secretory cells of the female reproductive tract. eLife 2, e00415.

Tabata, T., Schwartz, C., Gustavson, E., Ali, Z., and Kornberg, T.B. (1995). Creating a *Drosophila* wing de novo, the role of engrailed, and the compartment border hypothesis. Development 121, 3359–3369.

Terhzaz, S., O’connell, F.C., Pollock, V.P., Kean, L., Davies, S.A., Veenstra, J.A., and Dow, J.A. (1999). Isolation and characterization of a leucokinin-like peptide of *Drosophila melanogaster*. J Exp Biol 202, 3667–3676.

Tricoire, H., Battisti, V., Trannoy, S., Lasbleiz, C., Pret, A.M., and Monnier, V. (2009). The steroid hormone receptor EcR finely modulates *Drosophila* lifespan during adulthood in a sex-specific manner. Mech Ageing Dev 130, 547–552.

Vallejo, D.M., Juarez-Carreno, S., Bolivar, J., Morante, J., and Dominguez, M. (2015). A brain circuit that synchronizes growth and maturation revealed through Dilp8 binding to Lgr3. Science 350, aac6767.

Veenstra, J.A., Agricola, H.J., and Sellami, A. (2008). Regulatory peptides in fruit fly midgut. Cell Tissue Res 334, 499–516.

Yoshinari, Y., Ameku, T., Kondo, S., Tanimoto, H., Kuraishi, T., Shimada-Niwa, Y., and Niwa, R. (2020). Neuronal octopamine signaling regulates mating-induced germline stem cell proliferation in female *Drosophila melanogaster*. bioRxiv, 2020.2003.2020.999938.

Zandawala, M., Marley, R., Davies, S.A., and Nässel, D.R. (2017). Characterization of a set of abdominal neuroendocrine cells that regulate stress physiology using colocalized diuretic peptides in *Drosophila*. Cell Mol Life Sci.

Zinke, I., Kirchner, C., Chao, L.C., Tetzlaff, M.T., and Pankratz, M.J. (1999). Suppression of food intake and growth by amino acids in *Drosophila*: the role of pumpless, a fat body expressed gene with homology to vertebrate glycine cleavage system. Development 126, 5275–5284.

