## Supplemental figures for "*Drosophila* insulin-like peptide 8 (DILP8) in ovarian follicle cells regulates ovulation and metabolism"

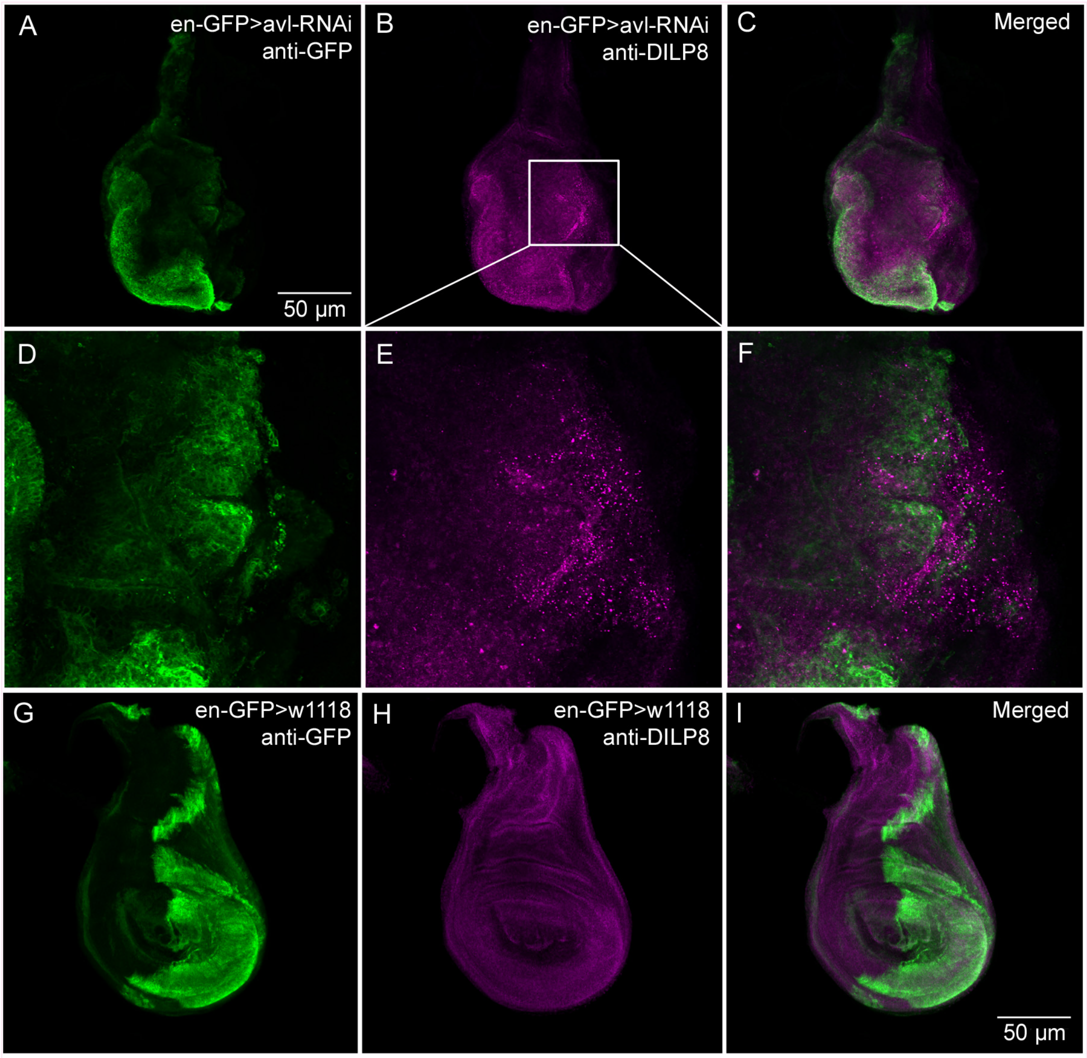


**Supplementary figure 1**. DILP8 immunoreactivity can be induced in wing imaginal discs by disrupting development. **A-C**. Using an *engrailed* (en) Gal4-driver (with GFP insertion) to knock down avalanche (en-GFP>avl-RNAi) we could induce DILP8 immunolabeling (framed area in B). GFP marks engrailed expression. **D-F.** Higher magnification of the area framed in B. **G-I.** Controls (en-GFP>$w^{1118}$) where on DILP8 immunolabeling is seen (**H**).

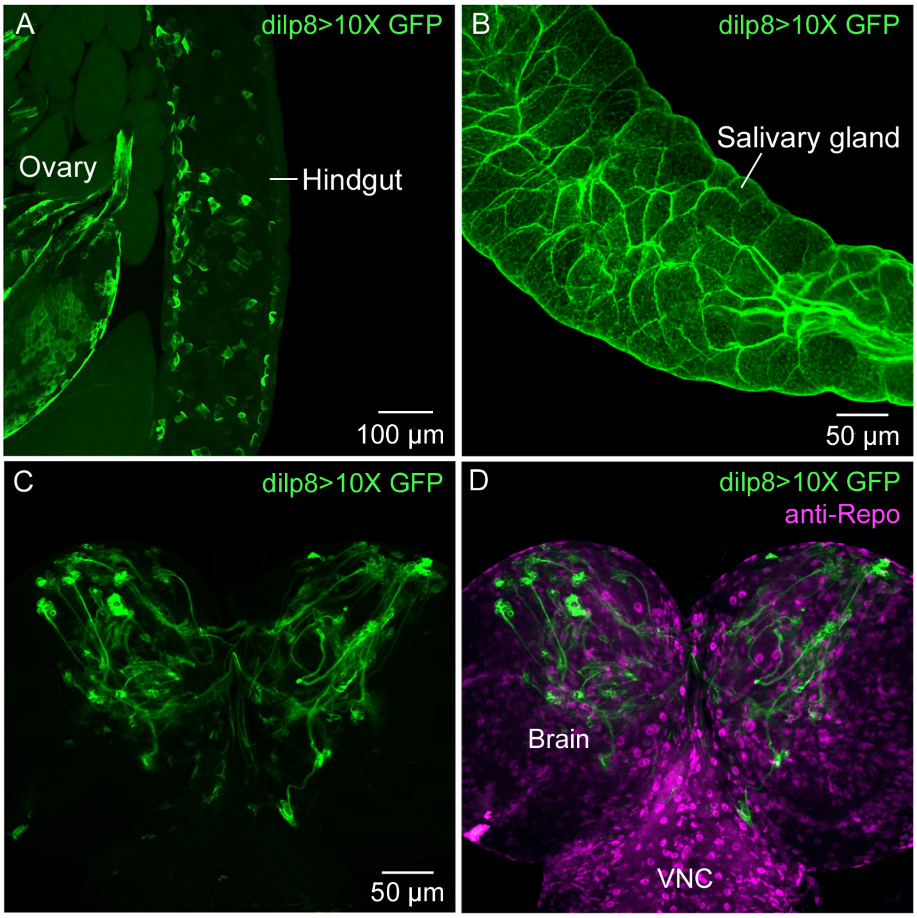


**Supplementary figure 2**. Further *dilp8* expression in tissues of adult and larval *Drosophila*. **A.** Using dilp8-Gal4 to drive 10X-GFP we could demonstrate expression in follicle cells of ovary and in cells of hindgut. **C.** Expression of *dilp8* in cells of salivary gland of adult fly. **C** and **D**. In third instar larvae neuronal progenitor cells of the brain express *dilp8*. In **D** glial cells are shown with repo-immunolabeling. VNC, ventral nerve cord.

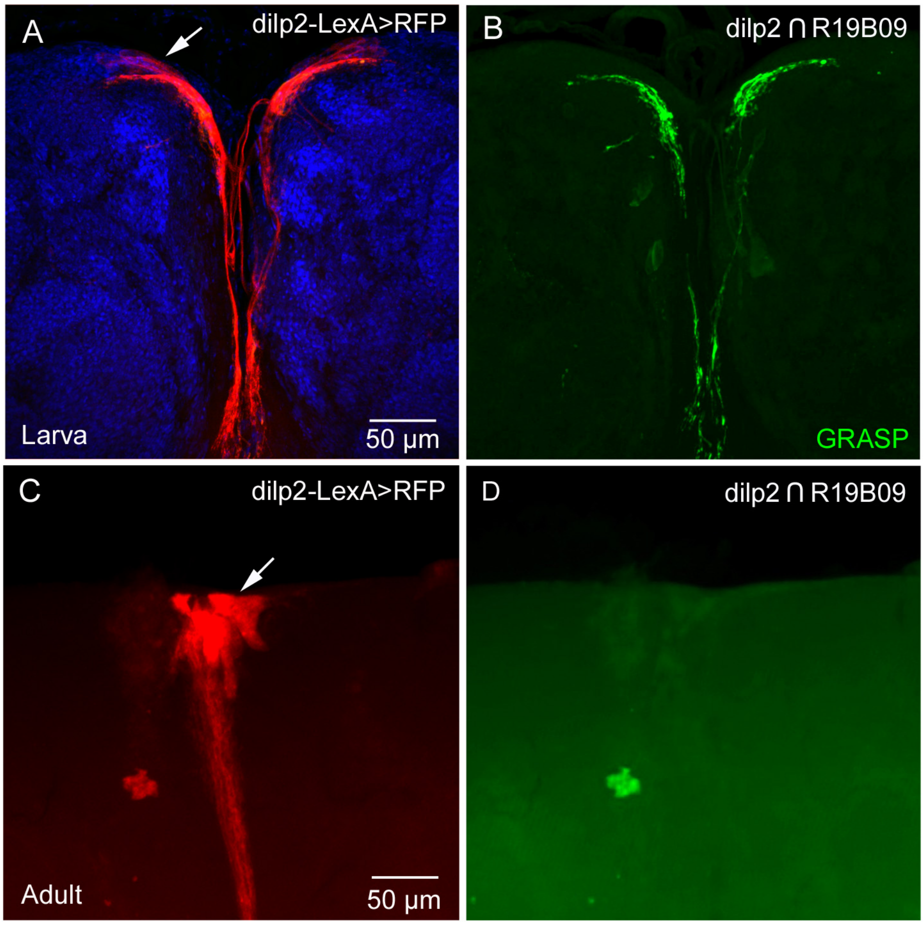


**Supplementary figure 3**. GFP-reconstitution across synaptic partners (GRASP) reveals contacts between Lgr3 neurons and IPCs. **A** and **B**. In the larval brain *dilp2*-LexA and R19B09-Gal4 generated GRASP (R19B09 Gal4>UAS-spGFP1-10; dilp2-lexA>LexAop-spGFP11; see methods) results in reconstituted GFP suggesting connections between IPCs and *Lgr3* expressing neurons (**B**). In A (and C) the red is red fluorescent protein (RFP) driven by *dilp2*-LexA; blue is DAPI labeling of nuclei. **C** and **D**. In adults GRASP with the same fly lines does not result in GFP, suggesting IPCs and Lgr3 neurons no longer form connections. Arrows in A and C points at cell bodies of IPCs (weakly labeled).

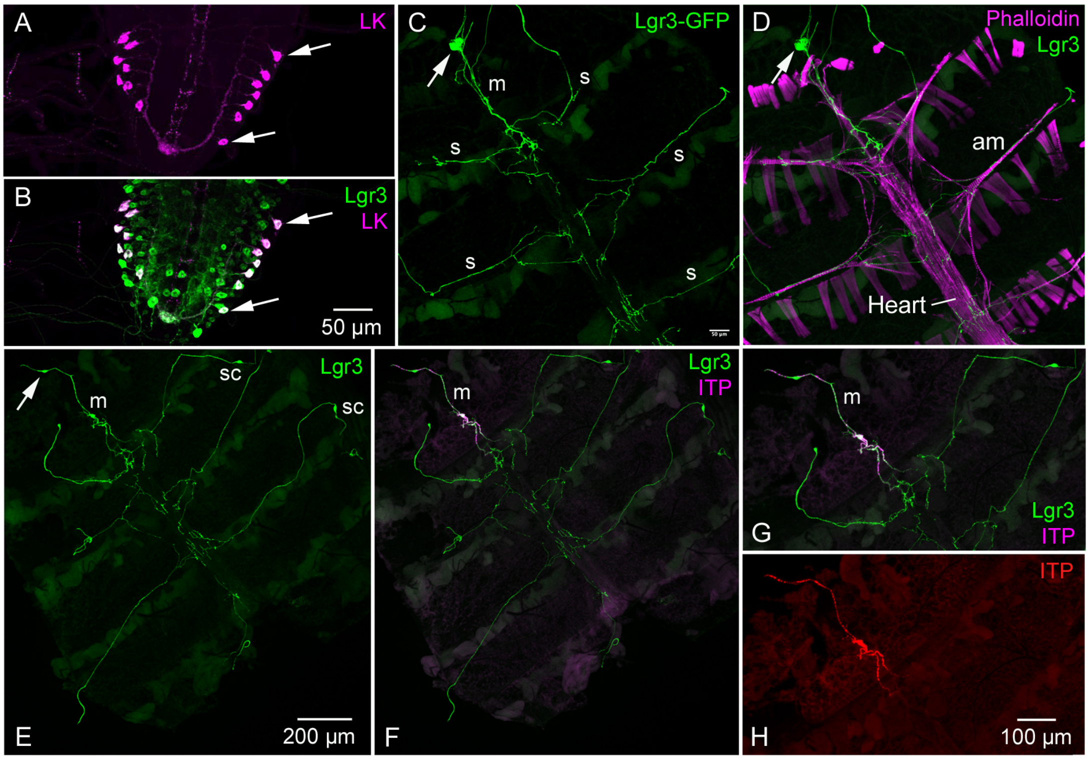


**Supplementary figure 4**. Lgr3 expression in larval CNS and adult peripheral neurons. **A** and **B**. In the larval abdominal neuromeres *Lgr3* is expressed in 7 pairs of leucokinin (LK)-producing neurosecretory cells (ABLKs) indicated by arrows. **C**-**H**. *Lgr3* is expressed in a small set of peripheral neurons. Some of these are lateral and segmental (s) and one pair is median (m), indicated by arrows in **C**-**E**). Both types of neurons innervate the dorsal aorta, or heart (see **D**). The segmental axons run along the alary muscles (am). The median *Lgr3* neurons express ion transport peptide (ITP) immunoreactivity (**F**-**H**). Note that the cell bodies are not clearly labeled with the ITP antiserum (**G** and **H**).

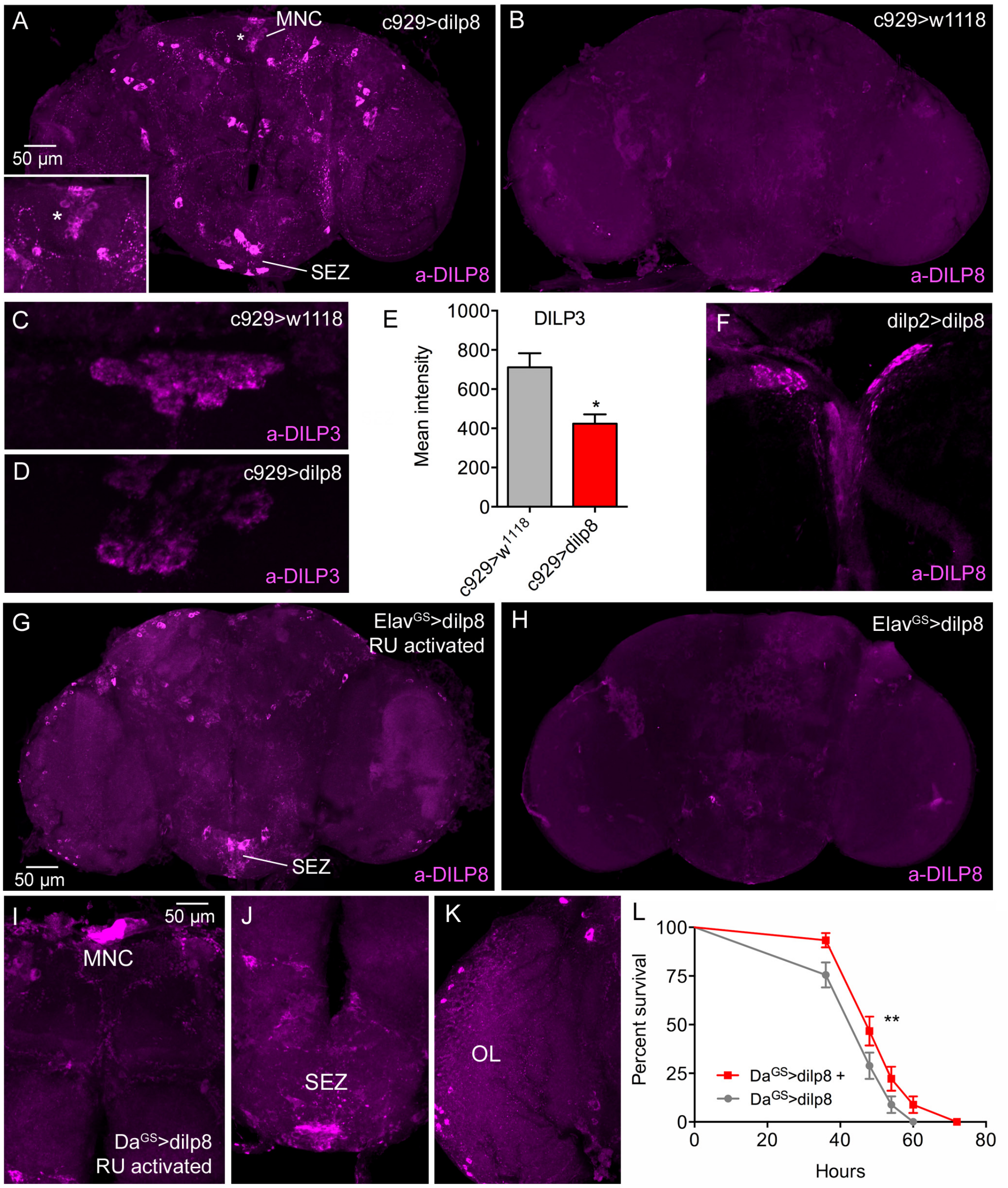


**Supplementary figure 5**. Overexpression of *dilp8* in different neurons results in DILP8 immunolabeling. **A**. The *c929*-Gal4>*dilp8* results in DILP8 immunolabeling in many peptidergic neurons, including median neurosecretory cells (MNC; see also enlarged at * in inset) and large neurons in the subesophageal zone (SEZ). **B.** In controls (c929>$w^{1118}$) no DILP8 immunolabeling can be detected. **C** and **D**. *c929*-Gal4>*dilp8* results in decreased DILP3 immunolabeling. **E.** Quantification of DILP3 labeling [5 flies used for *c929>dilp8* and 10 flies used for *c929>*$w^{1118}$ (*$p < 0.05$, unpaired Student's t-test)]. **F.** Ectopic expression of dilp8 in IPCs (dilp2>dilp8) results in DILP8 immunoreactivity. **G** and **H**. Conditional gene switch Gal4 expression (ElavGS>dilp8) induced in adult flies by feeding RU486 triggered DILP8 immunoreactivity in numerous neurons (**G**), and no labeling was seen without feeding RU486 (**H**). **I-K**. DILP8 immunolabeling in brain of $Da^{GS}$>*dilp8* flies fed RU486 as adults. DILP8 immunolabeling can be seen in median neurosecretory cells (MNC), neurons of subesophageal zone (SEZ) and optic lobe (OL). **L.** Starvation resistance of $Da^{GS}$>*dilp8* flies with RU486 activation (+) at adult stage increases compared to inactivated flies. 45 flies from three independent replicates were used [**$p < 0.01$, as assessed by log-rank (Mantel–Cox) test].
